# CACHE Challenge #1: targeting the WDR domain of LRRK2, a Parkinson’s Disease associated protein

**DOI:** 10.1101/2024.07.18.603797

**Authors:** Fengling Li, Suzanne Ackloo, Cheryl H. Arrowsmith, Fuqiang Ban, Christopher J. Barden, Hartmut Beck, Jan Beránek, Francois Berenger, Albina Bolotokova, Guillaume Bret, Marko Breznik, Emanuele Carosati, Irene Chau, Yu Chen, Artem Cherkasov, Dennis Della Corte, Katrin Denzinger, Aiping Dong, Sorin Draga, Ian Dunn, Kristina Edfeldt, Aled Edwards, Merveille Eguida, Paul Eisenhuth, Lukas Friedrich, Alexander Fuerll, Spencer S Gardiner, Francesco Gentile, Pegah Ghiabi, Elisa Gibson, Marta Glavatskikh, Christoph Gorgulla, Judith Guenther, Anders Gunnarsson, Filipp Gusev, Evgeny Gutkin, Levon Halabelian, Rachel J. Harding, Alexander Hillisch, Laurent Hoffer, Anders Hogner, Scott Houliston, John J Irwin, Olexandr Isayev, Aleksandra Ivanova, Austin J Jarrett, Jan H. Jensen, Dmitri Kireev, Julian Kleber, S. Benjamin Koby, David Koes, Ashutosh Kumar, Maria G. Kurnikova, Alina Kutlushina, Uta Lessel, Fabian Liessmann, Sijie Liu, Wei Lu, Jens Meiler, Akhila Mettu, Guzel Minibaeva, Rocco Moretti, Connor J Morris, Chamali Narangoda, Theresa Noonan, Leon Obendorf, Szymon Pach, Amit Pandit, Sumera Perveen, Gennady Poda, Pavel Polishchuk, Kristina Puls, Vera Pütter, Didier Rognan, Dylan Roskams-Edris, Christina Schindler, François Sindt, Vojtěch Spiwok, Casper Steinmann, Rick L. Stevens, Valerij Talagayev, Damon Tingey, Oanh Vu, W. Patrick Walters, Xiaowen Wang, Zhenyu Wang, Gerhard Wolber, Clemens Alexander Wolf, Lars Wortmann, Hong Zeng, Carlos A. Zepeda, Kam Y. J. Zhang, Jixian Zhang, Shuangjia Zheng, Matthieu Schapira

## Abstract

The CACHE challenges are a series of prospective benchmarking exercises meant to evaluate progress in the field of computational hit-finding. Here we report the results of the inaugural CACHE #1 challenge in which 23 computational teams each selected up to 100 commercially available compounds that they predicted would bind to the WDR domain of the Parkinson’s disease target LRRK2, a domain with no known ligand and only an apo structure in the PDB. The lack of known binding data and presumably low druggability of the target is a challenge to computational hit finding methods. Seventy-three of the 1955 procured molecules bound LRRK2 in an SPR assay with K_D_ lower than 150 μM and were advanced to a hit expansion phase where computational teams each selected up to 50 analogs each. Binding was observed in two orthogonal assays with affinities ranging from 18 to 140 μM for seven chemically diverse series. The seven successful computational workflows varied in their screening strategies and techniques. Three used molecular dynamics to produce a conformational ensemble of the targeted site, three included a fragment docking step, three implemented a generative design strategy and five used one or more deep learning steps. CACHE #1 reflects a highly exploratory phase in computational drug design where participants sometimes adopted strikingly diverging screening strategies. Machine-learning accelerated methods achieved similar results to brute force (e.g. exhaustive) docking. First-in-class, experimentally confirmed compounds were rare and weakly potent, indicating that recent advances are not sufficient to effectively address challenging targets.

## INTRODUCTION

The Critical Assessment of Computational Hit-finding Experiments (CACHE) is a triannual series of prospective benchmarking challenges in which computational chemistry experts are invited to select up to 100 compounds from commercial libraries that they predict bind to a pre-defined target. Compounds are purchased and binding to the protein is tested experimentally. Compounds of interest are then advanced to a hit expansion round where participants select up to 50 follow-up molecules for experimental testing. Based on both rounds, an independent committee composed of industry experts assesses the validity of the biophysical activity data of each series, the drug-likeness of the validated hits and their suitability as starting points for hit-to-lead optimization. Both the structures and bioactivity data serve to identify the best-performing computational methods, after which all data are publicly released on https://cache-challenge.org/. The goal of CACHE is to provide a unifying metric where a diverse array of virtual screening workflows are compared against the same protein target and evaluated using the same experimental assays and platform^1^.

The first CACHE challenge was focused on leucine rich repeat kinase 2 (LRRK2), the most mutated protein in familial Parkinson’s disease (PD). Mutations in the kinase domain of LRRK2 can increase its kinase activity and lead to pathogenic hallmarks associated with PD^2–4^. The kinase activity of LRRK2 has been an active area of drug discovery, but the first-generation LRRK2 kinase inhibitors have not demonstrated the therapeutic benefit hoped-for, which may be due to its scaffolding function^5^ or to the distinct conformational states stabilized by Type I and Type II inhibitors^6^. An alternative and so far overlooked strategy to inhibit pathogenic LRRK2 is to pharmacologically target its WD40 repeat (WDR) domain (LRRK2-WDR), which is juxtaposed to the kinase domain^7^ (Figure 1). A number of WDR domains are disease-associated and were shown to be druggable^8,9^. The WDR in LRRK2 domain may be important to recruit LRRK2 binding partners or for binding to tubulin. The WDR domain is also relevant to PD pathogenesis: a disease-linked mutation in the WDR domain is found at the interface of the LRRK2-WDR dimer, enhances LRRK2 kinase activity, and antagonizes dimerization^7^. Identifying compounds binding the LRRK2-WDR is a potentially novel approach to target this protein, though no ligand was reported to date. CACHE #1 participants were challenged to use the apo structure of LRRK2-WDR (PDB code 6DLO)^7^ to predict compounds occupying the central cavity of the donut shaped domain (Figure 1).

**Figure 1:**
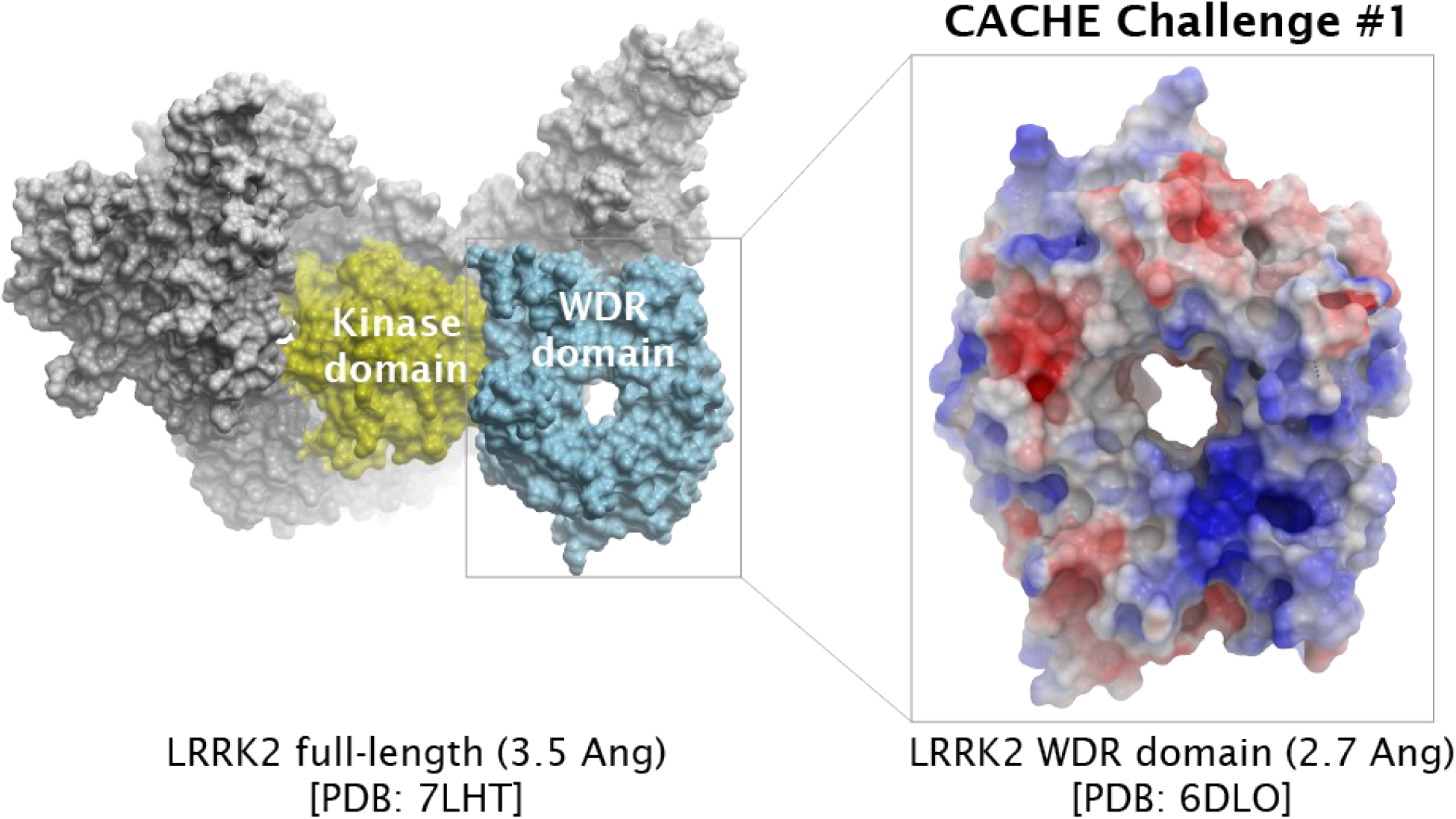
CACHE challenge #1: predicting ligands binding the central cavity of the LRRK2-WDR domain. Left: the kinase and WDR domains of LRRK2 are highlighted in the context of a LRRK2 monomer (PDB: 7LHT). Right: Electrostatic potential map of the LRRK2-WDR (blue: electropositive, red electronegative).

We provide here an overview of the first CACHE challenge, where 23 research teams from ten countries collectively predicted 1955 compounds targeting LRRK2-WDR. After a hit identification round (Round 1) followed by hit expansion (Round 2), seven chemical series predicted by seven participants produced convincing binding data in two orthogonal assays. These compounds are the first reported that target LRRK2-WDR and represent valuable chemical starting points for hit-to-lead optimization. Computational workflows were diverse and often included a step driven by deep learning. Hit rates were low and most compounds bound with an affinity above 50 μM, reflecting the challenges of structure-based virtual screening when only an apo form of the targeted binding pocket and no ligand is available.

## RESULTS

### Computational workflows were diverse

Participating teams were selected based on a double-blind peer review process of their applications. Each team was first asked to rate five applications after which an independent Applications Review Committee (Table S1) undertook a final evaluation. Twenty-five computational teams were selected, 23 of which completed the challenge. Participants remained anonymous until the final release of the data, at the end of the challenge, at which point they had the option to be de-anonymized. CACHE #1 participants deployed a highly diverse set of computational tools and workflows, reflecting different hit selection strategies summarized in Figure 2 and detailed in https://cache-challenge.org/challenges/predict-hits-for-the-wdr-domain-of-lrrk2/computational-methods. In one instance, Shuangjia Zheng at Shanghai Jiao Tong University (workflow WF1187) used a multi-scale and multi-task neural network pre-trained on ChEMBL and PubChem data as a one-step virtual screening workflow to produce the final compound selection, refined with physicochemical drug-likeness filters^10^. Pavel Polishchuk at Palacky University (WF1210) adopted a highly contrasting screening cascade composed of seven distinct steps, where he first used molecular dynamics (MD) to generate a conformational ensemble of the binding pocket to which fragments were docked, grown and fine-tuned by a genetic algorithm for de-novo ligand design; a consensus docking step refined with Molecular Mechanics generalized Born Surface Area (MM-GBSA) simulations was used to select the most promising ligands, commercial analogs of which (found by a fingerprint similarity search) were subjected to consensus docking followed by MM-GBSA to produce the final selection.

**Figure 2:**
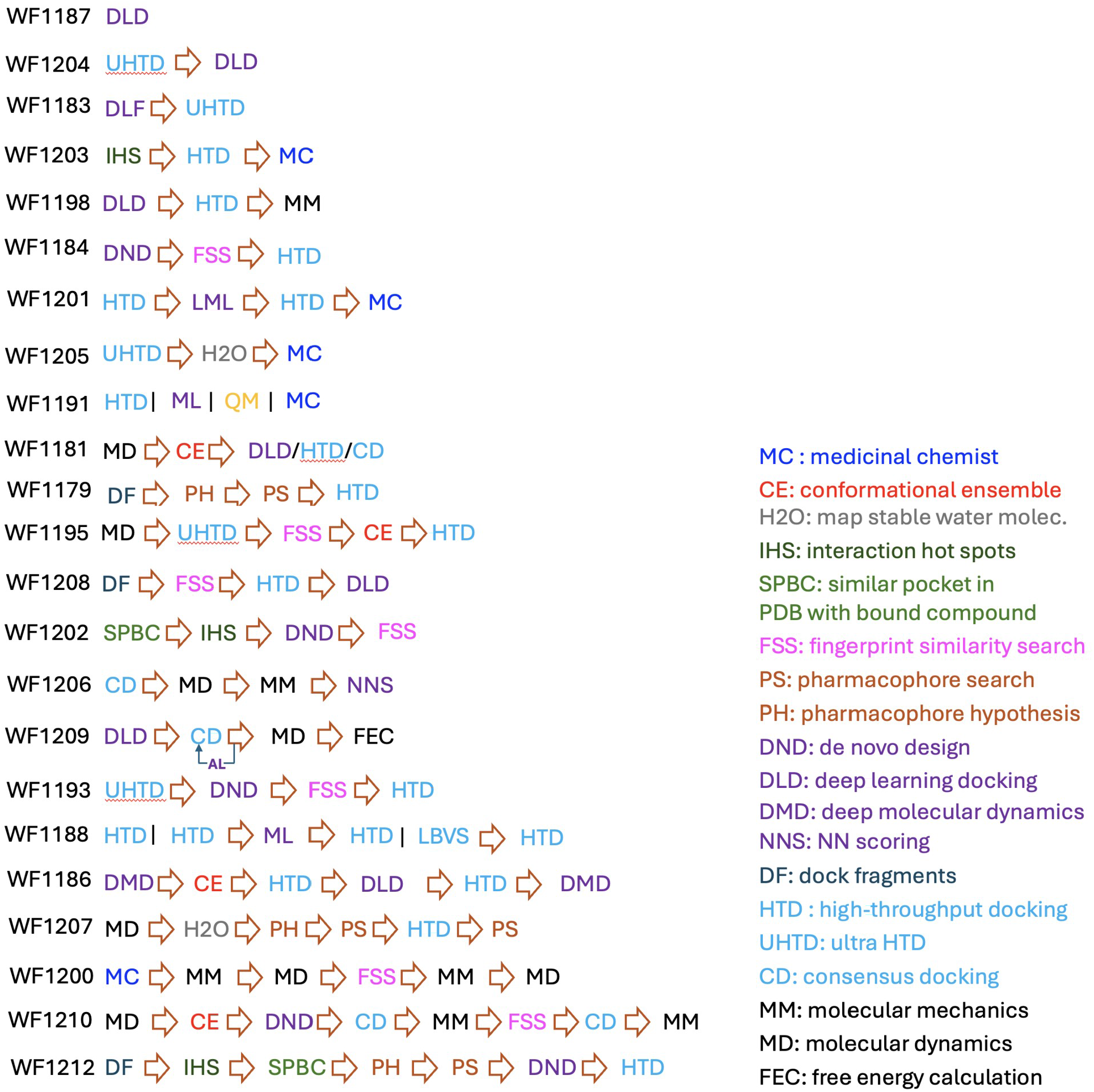
Computational cascades deployed in CACHE #1. Arrows denotes cascading steps. “|” denotes alternative methods tested in parallel. Each workflow had a maximum credit of 100 compounds, regardless of the number of methods tested.

Between these two extremes, which both ranked in the top 10 after experimental testing, screening cascades greatly varied in length and techniques (Figure 2). Physics-based docking was used in 19 workflows; 12 incorporated at least one deep learning screening step, including deep learning docking in eight. Fragment-based approaches were adopted in five, four used MD to generate a conformational ensemble of the binding site, and four included consensus docking.

### Selected compounds were drug-like and chemically diverse

After two months allocated to virtual screening, each of the 23 participants submitted a file of up to 100 compounds predicted to bind the central pocket of LRRK2-WDR and available from the Enamine REAL database (36 billion molecules at the time). Compounds had to satisfy three conditions: MW < 550 Da, cLogP < 5 and no reactive group. Participants were also encouraged to use badapple (https://datascience.unm.edu/badapple/)^11^ to filter promiscuous compounds, but this was not mandatory. Almost all compounds (1875) were procured from Enamine with a synthesis success rate of 93% and 80 were procured from MCULE, leading to 80 to 100 compounds for most participants, with a few exceptions, including participant 1183 who selected only 37 compounds. The distribution of physicochemical descriptors of the 1955 compounds selected by the 23 participants reflected overall drug-like molecules (Figure 3, Table S2).

**Figure 3.**
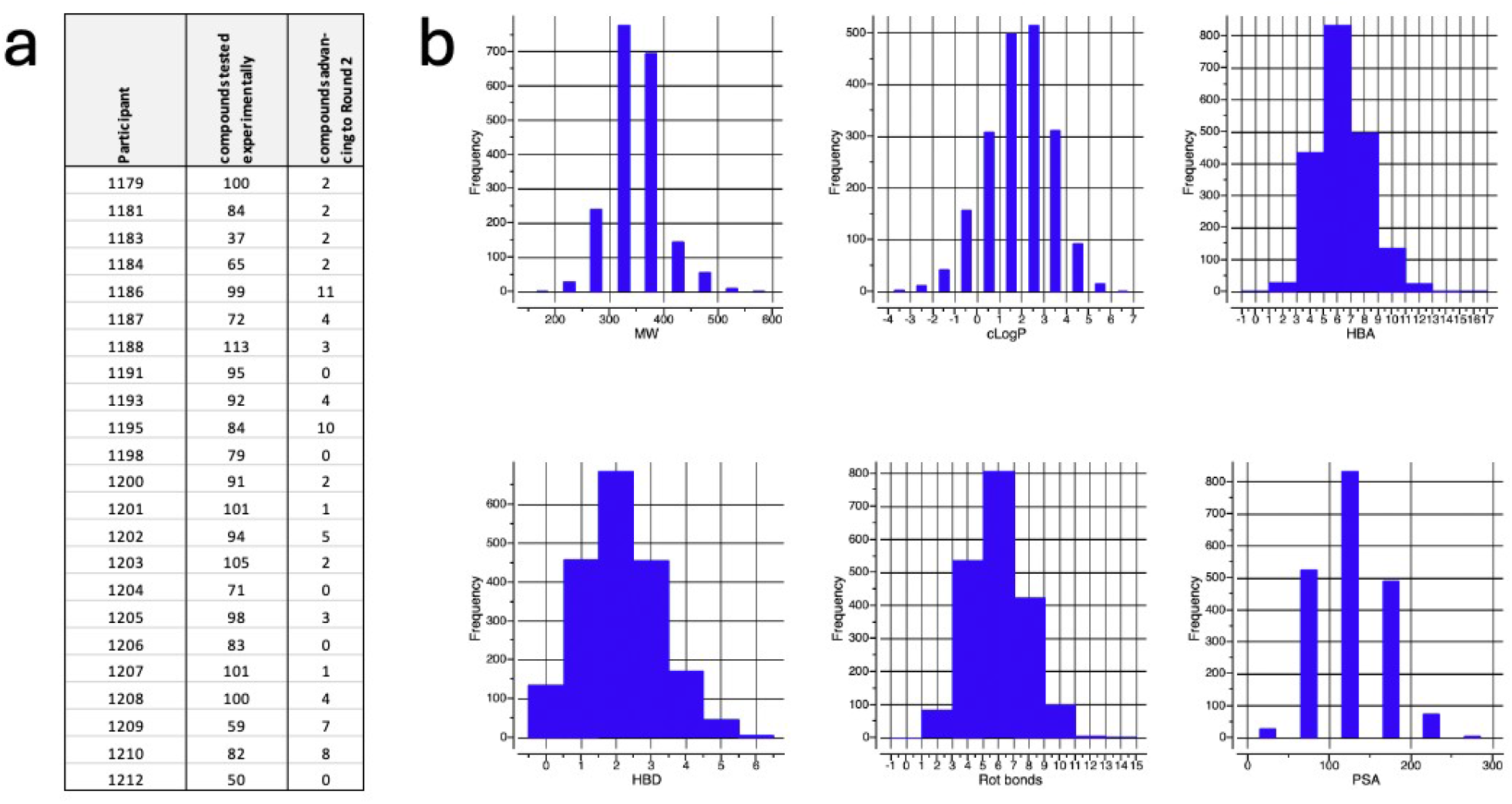
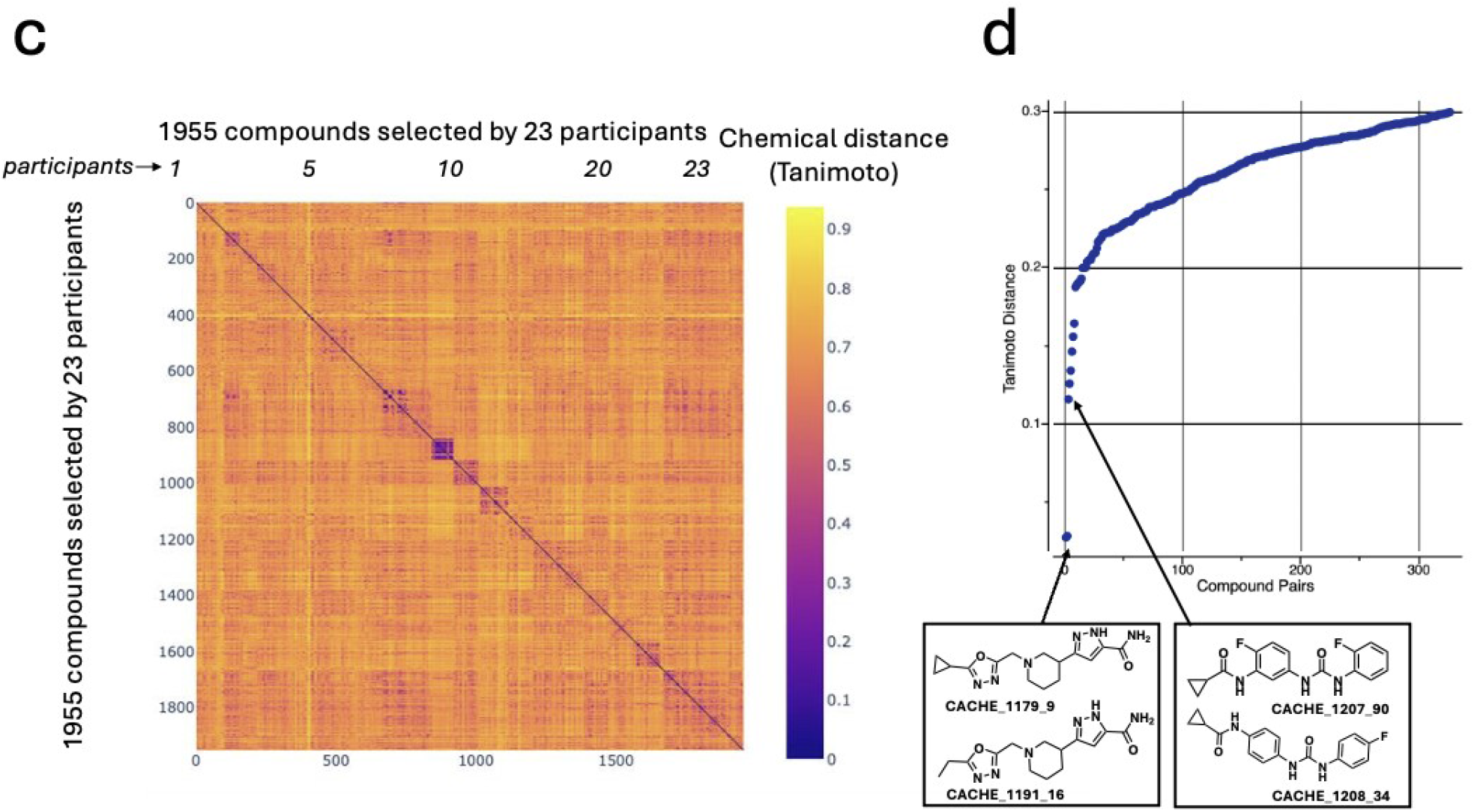
Drug-likeness and chemical diversity of selected compounds. a) Number of compounds procured for each participant in the hit identification phase (Round 1) and number advancing to hit expansion (Round 2). b) Molecular weight, calculated LogP, number of hydrogen bond acceptors /donors and rotatable bonds, and polar surface area distributions of the 1955 compounds. c) Pairwise Tanimoto distance matrix of all pairs of compounds. d) Tanimoto distance distribution of the 326 closest compound pairs (pairs of compounds selected by the same participant are not included).

The compounds were chemically diverse, with 1629 compounds out of 1955 having a Tanimoto distance greater than 0.3 with any compound selected by another participant (using 1536-bit fingerprints implemented in ICM, Molsoft LLC) (Figure 3c,d). Chemical diversity was also observed within selections from each participant, though some participants did select multiple chemically related compounds (dark squares along the diagonal in Figure 3c).

### Experimental testing of Round 1 compounds

Binding of the 1955 Round 1 compounds to LRRK2-WDR was tested independently at 50 μM and 100 μM compound concentrations in a surface plasmon resonance (SPR) assay (Figure 4a, Table S3). 440 compounds with a R/R_max_ binding ratio (measured versus expected response unit (RU)) above 50% (i.e. significant binding) and below 200% (i.e. limited signs of non-specific binding) in at least one of the two runs were evaluated in dose-response experiments. In total 73 compounds selected by 18 participants had a measurable dissociation constant K_D_ value better than 150 μM and greater than 30% binding (R/R_max_) (Table S4). To assess that the binding signal was on target, SPR was used to evaluate binding to an unrelated target, the first PWWP domain of the protein methyltransferase NSD2 (NSD2-PWWP1). Seventeen compounds bound NSD2-PWWP1 with K_D_ values ranging from 2 to 177 μM.

**Figure 4.**
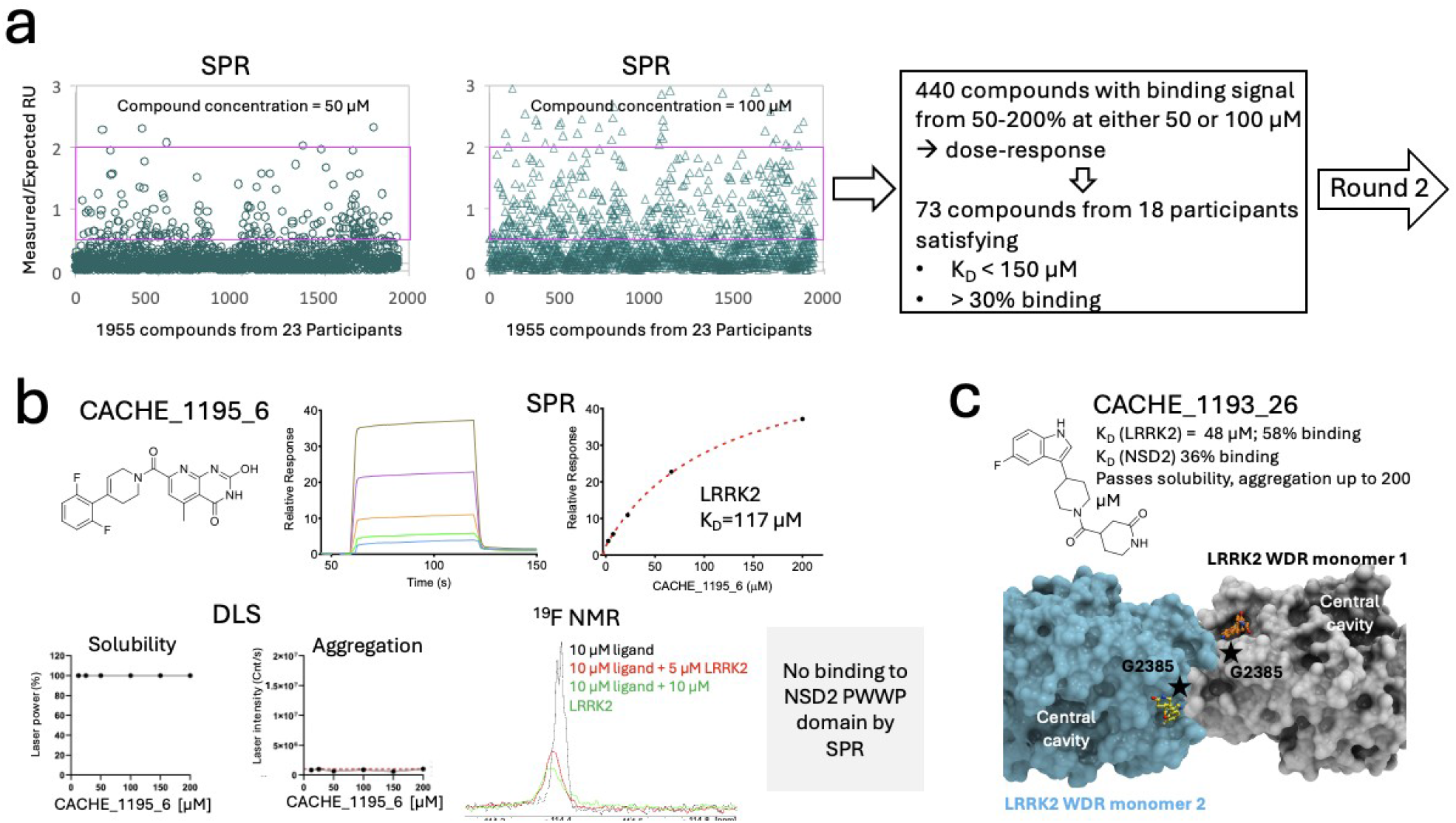
Experimental evaluation of CACHE Round 1 compounds. a) Binding to LRRK2 measured by SPR was used to advance compounds to Round 2. b) Experimental data beyond SPR was provided to better inform participants, including solubility and aggregation measured by DLS, binding by SPR to an unrelated protein (NSD2-PWWP1), and data from orthogonal binding assays (^19^F-NMR shown here). Data for compound CACHE_1195_6 is shown as an example. c) Crystal structure of CACHE_1193_26 bound at the interface of two LRRK2 monomers (PDB 9C61), distant from the targeted central cavity, at a site lined by G2385, recurrently mutated to Arg in PD patients.

Some of the 73 compounds showed signs of aggregation or poor solubility as measured by dynamic light scattering (DLS)^12^, none of the tested compounds showed clear signs of binding in differential scanning fluorimetry (DSF) or isothermal titration calorimetry (ITC) assays and only two compounds (CACHE_1195_6, SPR K_D_ 117 μM and CACHE_1210_69, K_D_ 117 μM) out of eleven tested bound to LRRK2-WDR in a ^19^F-NMR assay (Figure 4b). The only successful co-crystallization or crystal soaking attempt was with compound CACHE_1193_26 (SPR K_D_ 46 μM). The binding pose captured experimentally was not at the central cavity where the compound was docked, but at the interface of two LRRK2-WDR monomers (Figure 4c, Table S5). To test the hypothesis that the binding mode captured in the crystal structure might be induced by crystallization, we generated and purified a mutant form of the LRRK2 WDR domain in which the glycine residue lining the observed pocket was replaced with arginine (LRRK2-WDR G2385R), a space filling residue that is also found in PD patients^7^. Using SPR, we observed that the compound bound equally well to the two forms, strongly suggesting that the binding pocket observed in the crystal structure is distinct from the one exploited in solution.

Some of the 73 SPR hits showed suboptimal behavior in solution, including 37 compounds with signs of poor solubility by DLS, some of which also produced a binding signal against the anti-target NSD2-PWWP1, and almost none were confirmed with an orthogonal biophysical assay. In spite of these red flags, we decided to advance all 73 compounds of interest to the hit expansion stage to avoid false negatives, which we discuss later.

### Selection and experimental testing of Round 2 compounds

While the focus of Round 1 was to avoid false negatives, Round 2 was equally focused on avoiding false positives. Here, participants selected up to 50 commercially available analogs of their compounds experimentally identified in Round 1. The aim was to generate structure activity relationship (SAR) to build confidence that binding signals were not artifacts from the assay or driven by other irrelevant factors such as aggregation. A total of 714 selected compounds were tested experimentally, representing 23 to 49 compounds per participant and up to 43 analogs per parent molecule (Table 1, Table S6). In total, analogs were tested for 42 of the 73 compounds of interest identified in Round 1 as participants focused on their more promising hit candidates and analogs were not always commercially available.

**Table 1:**
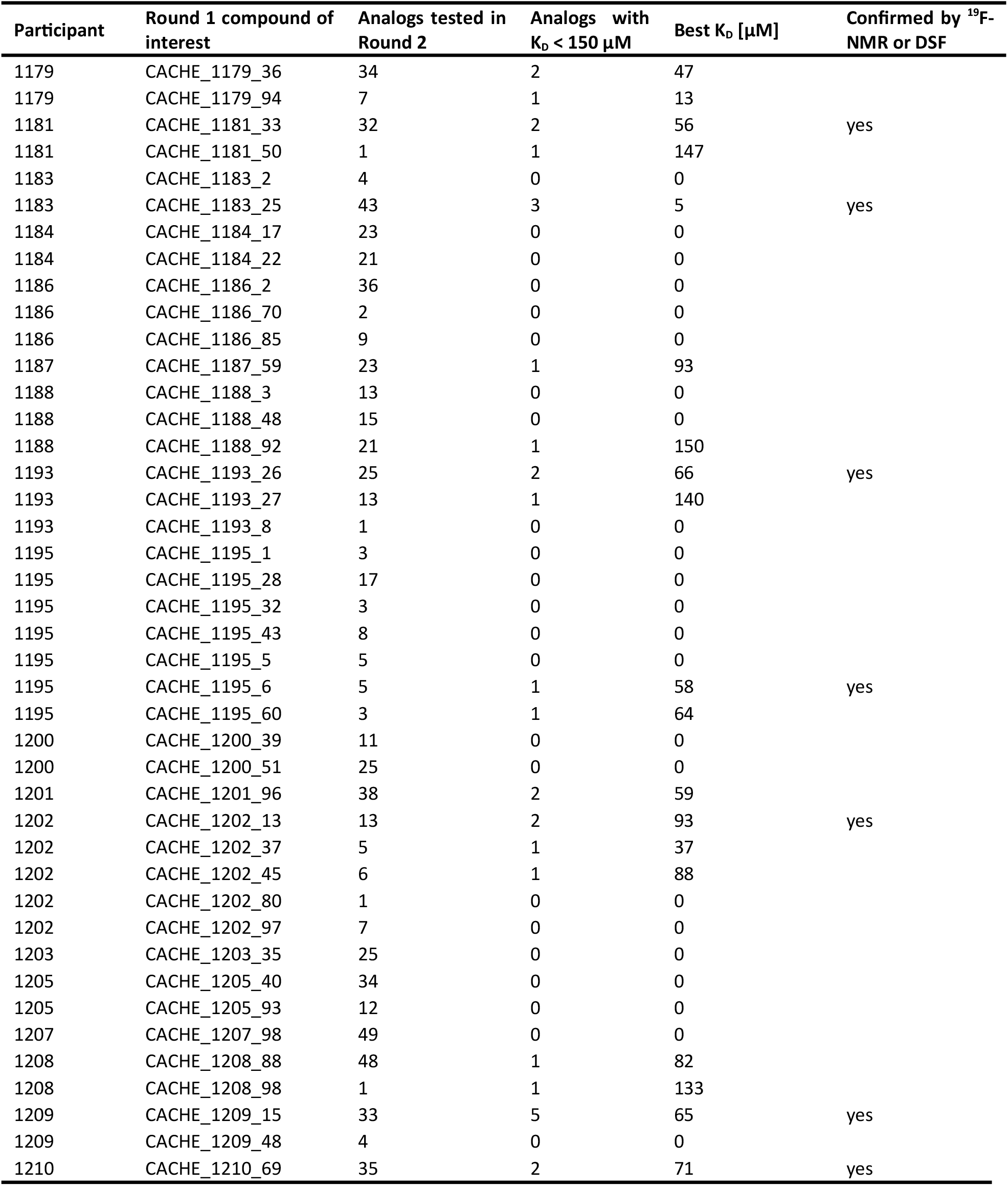
Summary of Round 2 experimental results.

| Participant | Round 1 compound of interest | Analog tested in Round 2 | Analog with $K_D < 150 \mu M$ | Best $K_D [\mu M]$ | Confirmed by $^{19}F$-NMR or DSF |
| --- | --- | --- | --- | --- | --- |
| 1179 | CACHE_1179_36 | 34 | 2 | 47 |  |
| 1179 | CACHE_1179_94 | 7 | 1 | 13 |  |
| 1181 | CACHE_1181_33 | 32 | 2 | 56 | yes |
| 1181 | CACHE_1181_50 | 1 | 1 | 147 |  |
| 1183 | CACHE_1183_2 | 4 | 0 | 0 |  |
| 1183 | CACHE_1183_25 | 43 | 3 | 5 | yes |
| 1184 | CACHE_1184_17 | 23 | 0 | 0 |  |
| 1184 | CACHE_1184_22 | 21 | 0 | 0 |  |
| 1186 | CACHE_1186_2 | 36 | 0 | 0 |  |
| 1186 | CACHE_1186_70 | 2 | 0 | 0 |  |
| 1186 | CACHE_1186_85 | 9 | 0 | 0 |  |
| 1187 | CACHE_1187_59 | 23 | 1 | 93 |  |
| 1188 | CACHE_1188_3 | 13 | 0 | 0 |  |
| 1188 | CACHE_1188_48 | 15 | 0 | 0 |  |
| 1188 | CACHE_1188_92 | 21 | 1 | 150 |  |
| 1193 | CACHE_1193_26 | 25 | 2 | 66 | yes |
| 1193 | CACHE_1193_27 | 13 | 1 | 140 |  |
| 1193 | CACHE_1193_8 | 1 | 0 | 0 |  |
| 1195 | CACHE_1195_1 | 3 | 0 | 0 |  |
| 1195 | CACHE_1195_28 | 17 | 0 | 0 |  |
| 1195 | CACHE_1195_32 | 3 | 0 | 0 |  |
| 1195 | CACHE_1195_43 | 8 | 0 | 0 |  |
| 1195 | CACHE_1195_5 | 5 | 0 | 0 |  |
| 1195 | CACHE_1195_6 | 5 | 1 | 58 | yes |
| 1195 | CACHE_1195_60 | 3 | 1 | 64 |  |
| 1200 | CACHE_1200_39 | 11 | 0 | 0 |  |
| 1200 | CACHE_1200_51 | 25 | 0 | 0 |  |
| 1201 | CACHE_1201_96 | 38 | 2 | 59 |  |
| 1202 | CACHE_1202_13 | 13 | 2 | 93 | yes |
| 1202 | CACHE_1202_37 | 5 | 1 | 37 |  |
| 1202 | CACHE_1202_45 | 6 | 1 | 88 |  |
| 1202 | CACHE_1202_80 | 1 | 0 | 0 |  |
| 1202 | CACHE_1202_97 | 7 | 0 | 0 |  |
| 1203 | CACHE_1203_35 | 25 | 0 | 0 |  |
| 1205 | CACHE_1205_40 | 34 | 0 | 0 |  |
| 1205 | CACHE_1205_93 | 12 | 0 | 0 |  |
| 1207 | CACHE_1207_98 | 49 | 0 | 0 |  |
| 1208 | CACHE_1208_88 | 48 | 1 | 82 |  |
| 1208 | CACHE_1208_98 | 1 | 1 | 133 |  |
| 1209 | CACHE_1209_15 | 33 | 5 | 65 | yes |
| 1209 | CACHE_1209_48 | 4 | 0 | 0 |  |
| 1210 | CACHE_1210_69 | 35 | 2 | 71 | yes |

As in Round 1, SPR was the primary assay. Sixty-one compounds had a measurable K_D_ value (8.5% hit-rate) with acceptable SPR parameters (maximum binding signal (Rmax) > 30% of the expected signal, T(K_D_) > 1 and Chi^2^ < 10% Rmax), 31 of which had a K_D_ < 150 μM (Table S3). ^19^F-NMR and DSF assays were used to orthogonally confirm SPR hits. All data were evaluated by an independent Hit Evaluation Committee composed of industry experts (Table S1). Overall, seven chemical series were convincingly confirmed with two orthogonal assays (Tables 1, S3, S7, Figure 5). These are the first reported molecules targeting LRRK2-WDR. Interestingly, some compounds of interest that displayed significant liability in Round 1 produced convincing chemical series in Round 2 (Figure 5). For instance, CACHE_1181_33 (K_D_ value of 123 μM) showed signs of insolubility and aggregation at 200 μM as measured by DLS, but its fluorinated analog, CACHE-HO_1181_24 (K_D_ value of 56 μM), was soluble, did not aggregate at 200 μM and showed a clear binding signal by ^19^F-NMR. This supports the decision to advance non-convincing compounds of interest from Round 1 to Round 2.

**Figure 5:**
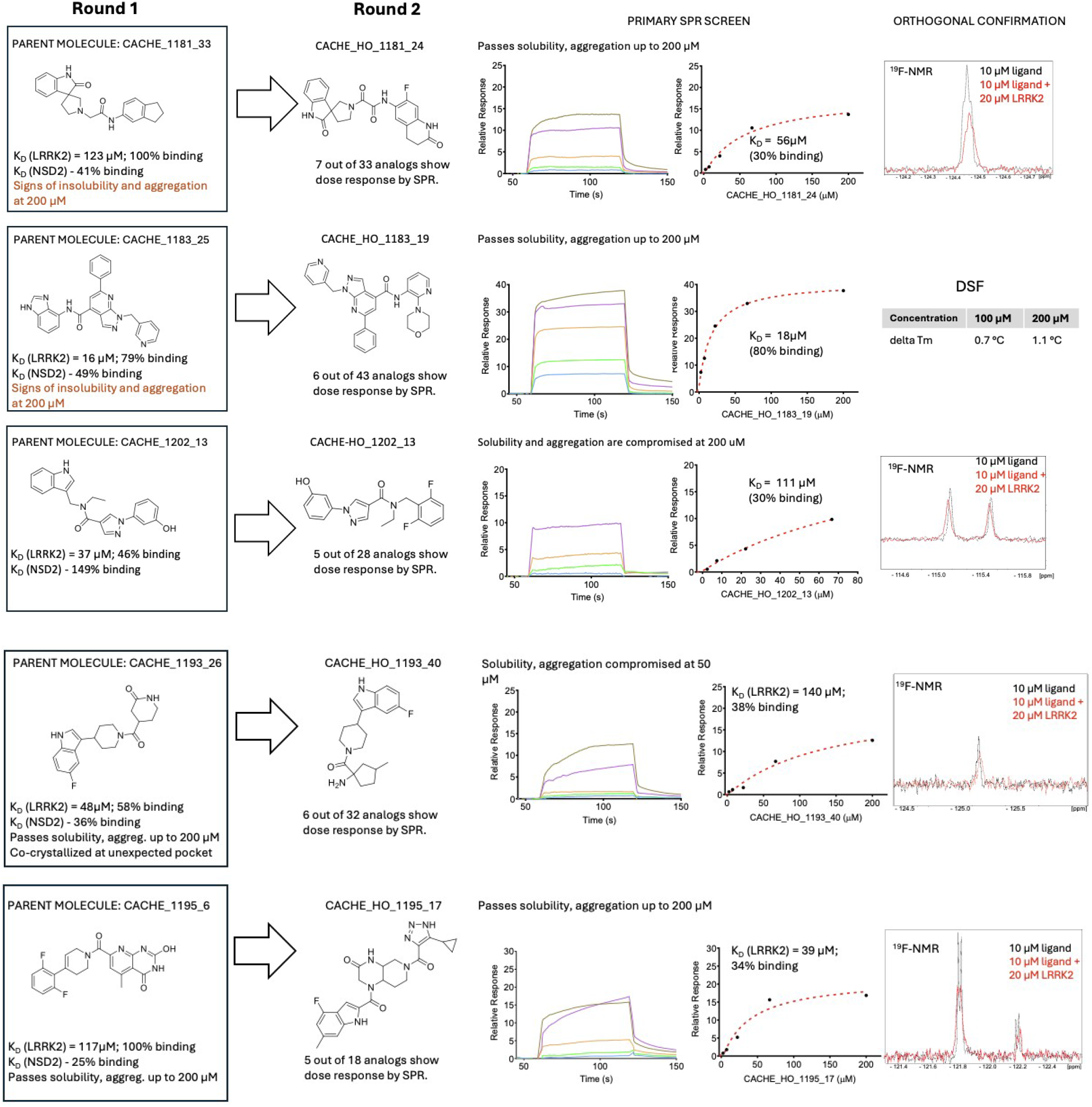
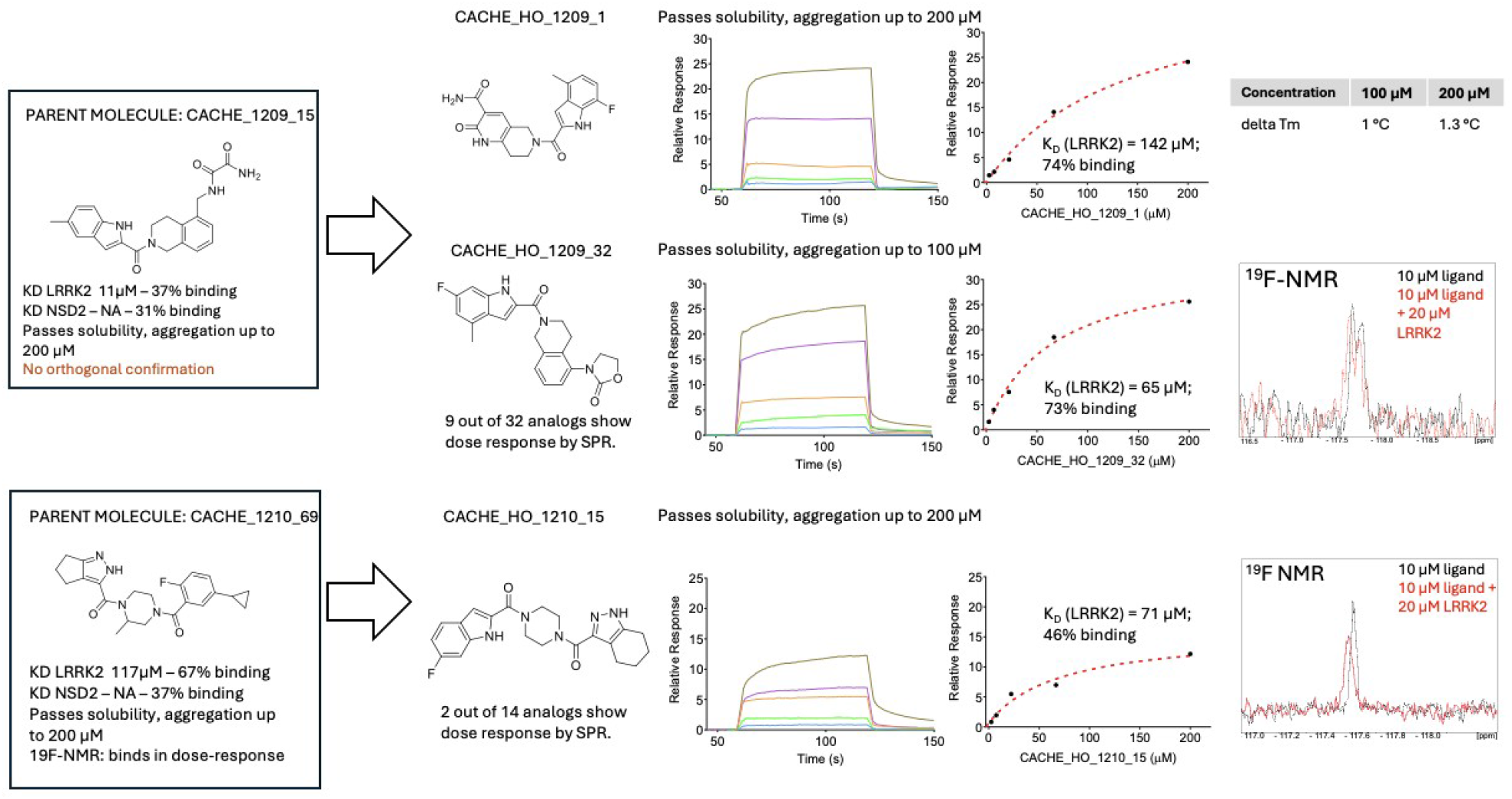
Top seven chemical series identified in Round 2. Activity of the parent molecules and experimental data from Round 2 analogs are shown, including SPR sensorgrams, ^19^F-NMR spectra and thermal shifts from DSF. Computational workflow IDs are encoded into compound names.

Nevertheless, we do not discount the possibility that some of the chemical series that were not validated by ^19^F-NMR (for lack of fluorinated compound) or by DSF (possibly due to distinct mode of binding) in Round 2 are still valid LRRK2-WDR ligands. A corollary is that CACHE should rather be used as a mechanism to highlight computational workflows that are performing well rather than those that are performing poorly. With this in mind, we next analyzed common and distinct features and design strategies adopted by the best performing CACHE participants.

### Emerging trends from the seven best performing computational workflows

Superimposing the docked poses of some of the top hits reveals that, while all were predicted to occupy the central channel of the LRRK2-WDR domain, there is no significant overlap in the predicted network of interactions with the protein, reflecting the open-ended and challenging nature of this binding site for structure-based drug design (Figure 6).

**Figure 6:**
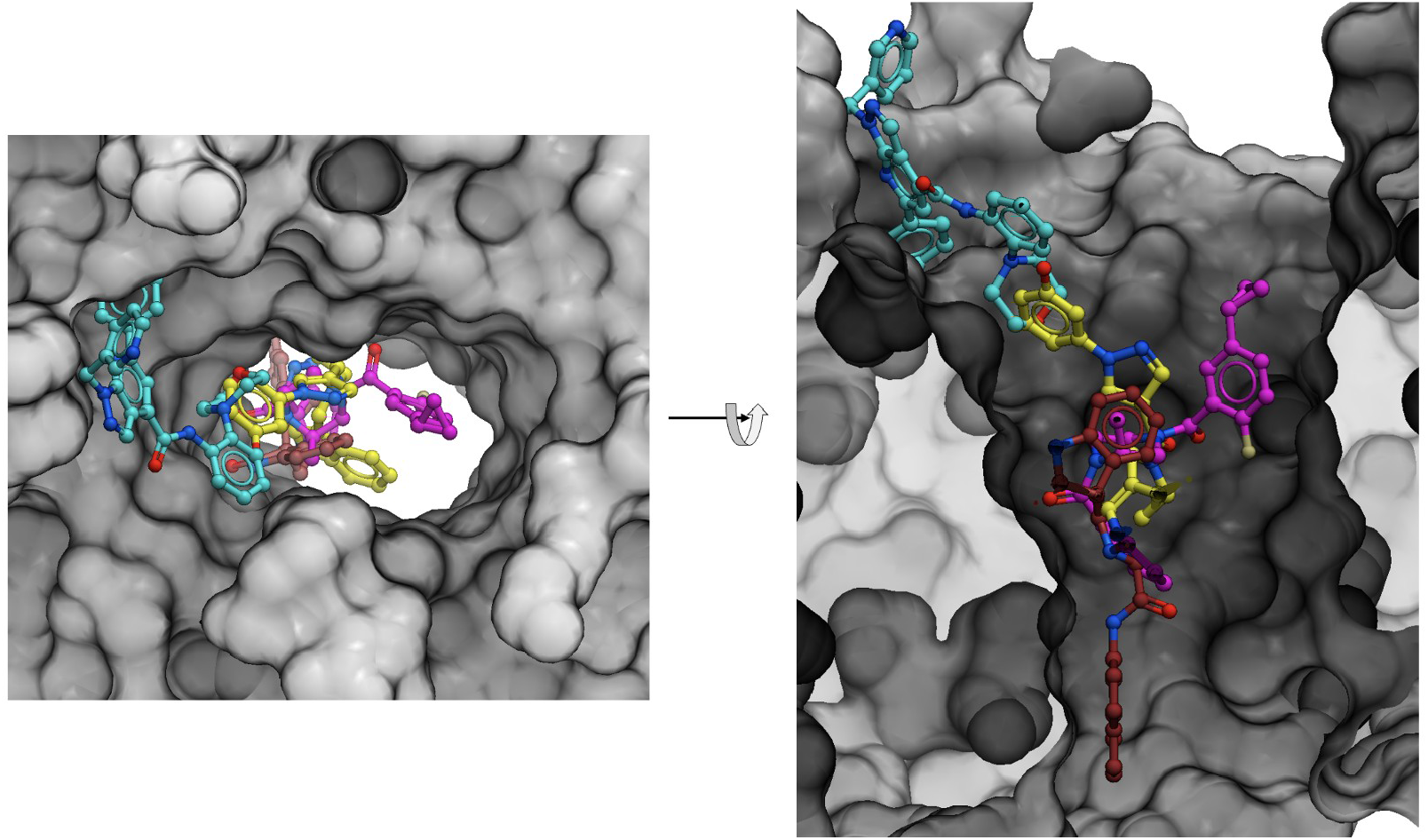
Docked poses of experimental hits. Four compounds are shown: CACHE_1183_13 (light blue), CACHE_1202_13 (yellow), CACHE_1210_69 (pink), CACHE_1181_33 (maroon). Top scoring poses for each ligand are shown. Computational workflows are included in the compound names and summarized in Figure 2.

The seven computational workflows that did produce a chemical series experimentally confirmed in two independent assays and had the best scores from the Hit Evaluation Committee were highly diverse (Figure 7, detailed description in https://cache-challenge.org/challenges/predict-hits-for-the-wdr-domain-of-lrrk2/computational-methods), but a few recurring trends and strategies were apparent.

**Figure 7:**
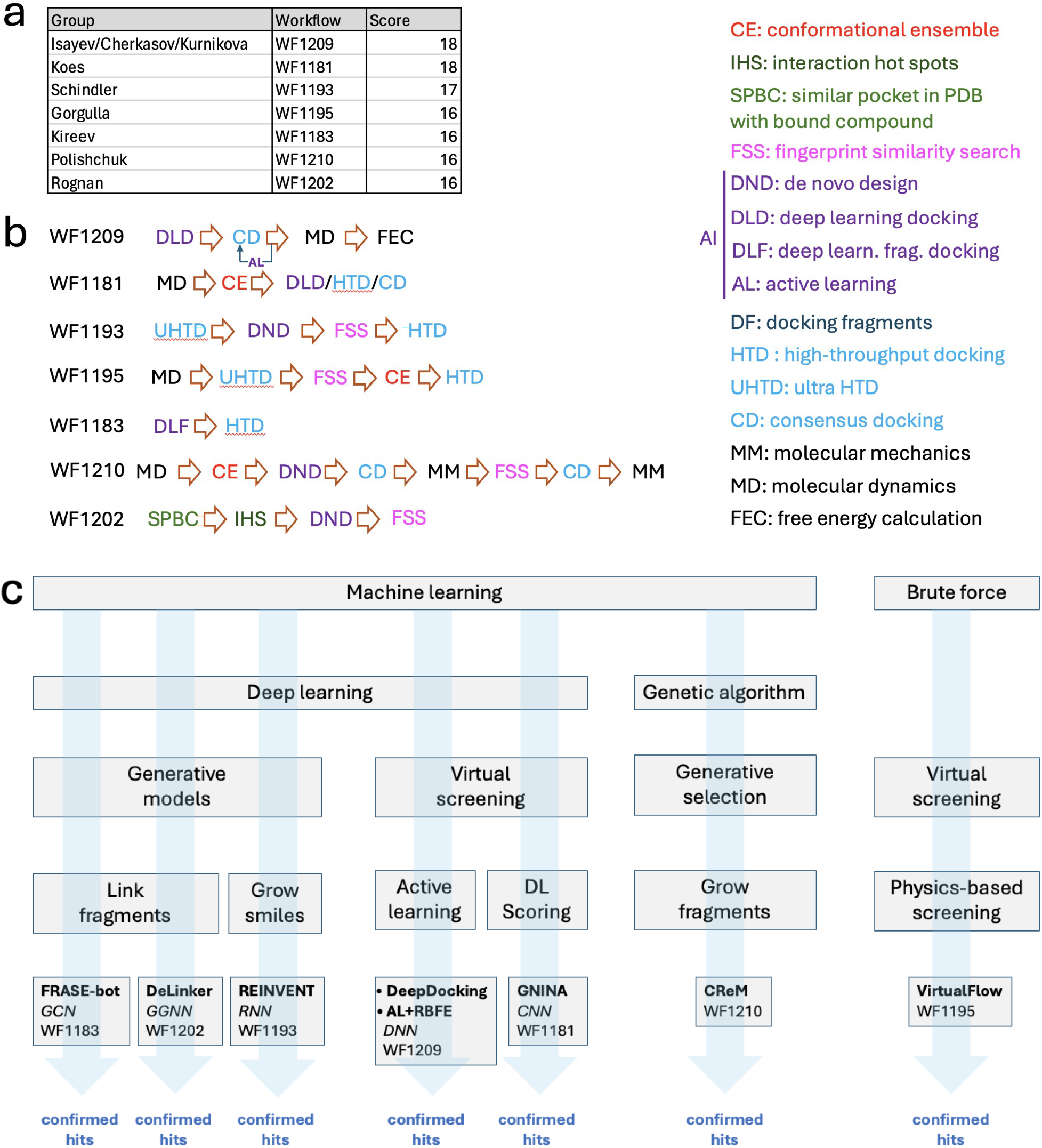
Overview of the best performing computational workflows. Details are provided for the seven computational workflows that had a chemical series experimentally confirmed with two binding assays. a) Team leads, workflow IDs and aggregated scores from the hit evaluation committee (Table S1 – Each committee member gave a score from 0 to 5 to each workflow based on the experimental data). b) Schematics representation of the computational steps for each workflow. c) Classification of the workflows based on some of their distinct features. Defining computational tools are outlined in bold. Neural network architectures are shown in italic. CNN: convolutional neural network; DNN: deep neural network; GCN: graph convolutional network; GGNN: gated graph neural network; RNN: recurrent neural network. DL: deep learning.

First, all workflows but one included one or more ML steps and five of these used some element of deep learning. Workflow 1181 (WF1181) adopted a physics-based high-throughput docking strategy complemented with a 3D convolutional neural network (CNN) scoring function implemented in GNINA^13^ to select compounds; WF1209 used DeepDocking^14^ where a deep neural network (DNN) predicts docking scores to rapidly screen an ultra-large library, followed by more refined active learning selection cycles where free energy calculation data was used to train an ML model. WF1193 used Glide docking scores (Schrödinger, New York) to train REINVENT, a recurrent neural network (RNN) and transformer-based ML model^15,16^ that generated de novo ligand candidates. WF1183 and WF1202 decomposed the binding site into local protein microenvironments with ligand-occupied structural homologs in the PDB; WF1183 used FRASE-bot^17^ where a graph convolutional network (GCN) distills an optimal feature vector from the protein-ligand interaction graphs to select the best fragments. Commercial molecules overlapping fragment pairs were then docked and ranked. WF1202 applied the POEM screening cascade where fragments positioned with a point cloud pocket registration system are linked with DeLinker, a generative ML model with a multimodal encoder-decoder set-up based on a standard gated graph neural network (GGNN)^18–20^. These successful strategies used deep learning to accelerate classical methods (WF1209), to predict affinity (WF1181), or to generate new molecules (WF1183, WF1193, WF1202). Finally, WF1210 did not use a neural network architecture but used CReM^21^ to grow previously docked fragments, followed by fine-tuning using a genetic algorithm. Workflow 1195 (WF1195) used VirtualFlow^22^ to deploy conventional physics-based virtual screening tools across tens of thousands of CPUs to efficiently dock an ultra-large chemical library.

Three of the workflows (WF1183, WF1202 and WF1210) used fragment-based approaches. Three used some form of de novo generative method to invent molecules customized for the target site (WF1193, WF1202, WF1210) followed by fingerprint similarity search to identify commercially available chemical analogs. As no ligand was reported at the outset of this challenge, available structures of the binding pocket were in the apo state, which is typically challenging for ligand binding and virtual screening. Three design strategies included molecular dynamics simulations to generate a conformational ensemble of the target site against which compounds were docked (WF1181, WF1195, WF1210).

Together, these results demonstrate that multiple design strategies and technical tools can successfully drive the structure-based discovery of pioneer ligands for an unprecedented target. Significant differences were also observed in the amount of computational resources used (Table S8). Two of the seven best performing workflows exclusively deployed conventional computational tools and methodologies that have been in use for decades (WF1195, WF1210), with results equivalent to those obtained with deep learning driven screening cascades, demonstrating that advanced neural network architectures did not lead to a breakthrough in this challenge.

## Discussion

Unlike previous computational challenges, where participants were asked to predict pre-generated experimental data blinded to them, CACHE is the first benchmarking challenge where computational predictions are experimentally tested prospectively. A new CACHE challenge is launched every four months, and for each CACHE target, suitable assays are used to confirm predicted hits. In this first iteration, computational workflows from seven independent teams used the apo structure of LRRK2-WDR to predict ligands that were subsequently confirmed experimentally. A number of lessons can be drawn from CACHE #1, both on the experimental and the in-silico aspects of the challenge. First, a major challenge faced by the experimental team was the overall poor solubility of the predicted molecules. Virtual screening can achieve high hit rates and produce potent molecules in the hands of seasoned experts when the structural chemistry of the target or target-class is well understood and known ligands are available to identify favorable pocket conformation(s), define interaction hotspots, and validate docking protocols^23–25^. Conversely, hit rates are typically low or null and compounds are weak when computationally screening underexplored proteins with no known ligand, as was the case here. Predicted molecules therefore must be tested at high concentration (up to 200 μM in this challenge), where they all too often tend to precipitate or aggregate. Indeed, 53% of molecules tested by DLS in Round 1 were not 100% soluble at 200 μM in the SPR buffer minus detergent. To the best of our knowledge, solubility prediction, when not trained on a given chemical series, is typically not highly reliable. Nevertheless, a mechanism to filter-out poorly soluble compounds before they are procured and tested would improve the screening process. It would also limit uncertainties associated with the nomination of compounds of interest showing weak activity and poor solubility.

Here, we made the choice in Round 1 not to filter-out compounds that behaved poorly in solution, for instance due to low solubility. Indeed, unlike a typical drug discovery project, the CACHE experimental team does not have the luxury of overlooking second-tier hit candidates, as it is important to avoid designating as unsuccessful a computational pipeline that may have produced structurally valid molecules. This necessity leads to the advancement of dubious molecules to Round 2, where the focus is to identify convincing hits and clearly successful computational pipelines, that may (and in effect, sometimes did) stem from problematic compounds of interest from Round 1 (e.g.: Figure 5, WF1181, WF1183).

While this challenge was successful in providing a unifying metric to compare computational screening pipelines and highlight successful ones, we also see areas for future improvement. First, while we were able to clearly identify computational methods that did well, we cannot say with absolute certainty that other methods did not. Indeed, several of the workflows that did not rank in the top seven produced chemically related hits by SPR which we were not able to confirm either by DSF or by ^19^F-NMR. We cannot discount the possibility that some of these hits were valid but did not produce a detectable binding signal in the orthogonal assay(s). Second, in the white paper detailing the scope and operational set-up of CACHE challenges^1^, we planned a step where all participants blindly screened a library composed of the merged collection of all compounds predicted in Round 1 to compare all methods against the same library. Because few hits were confirmed experimentally and fewer of these (two or less) were predicted by each participant in this exercise, this data did not allow for a statistically significant analysis and this evaluation step was dismissed. Third, structure-based virtual screening is not an exact science and the same computational workflow may succeed in someone’s hands and fail in someone else’s. This may not be critical if the goal is to identify partners (i.e. a successful combination of team and technology) to engage with in drug discovery projects, but it can be seen as a drawback when evaluating scientific methodologies. One opinion, which is related to point number one above, is that CACHE remains a valuable metric to identify successful workflows: the experimental validation of active molecules necessarily implies that the workflow produced valid hits and that humans picked some of them. Conversely, if a pipeline fails to produce valid molecules, the most expert computational or medicinal chemists will be hard pressed to use their intuition to pick active compounds. An option to eliminate the human factor would be for participants to submit a containerized version of their pipelines instead of a collection of molecules. CACHE organizers would then blindly run the methods and select compounds. A similar process was for instance implemented in the CELPP challenges^26^.

## Conclusion

The first iteration of the CACHE challenges closely followed a process precisely defined by an assembly of stakeholders in computational hit finding from the public and private sectors^1^. Despite expected and unforeseen limitations encountered along the road, CACHE #1 highlighted successful computational pipelines in structure-based virtual screening and produced experimentally confirmed ligands with an entirely novel structural mechanism for LRRK2, an important PD target, that may serve as a starting point to interrogate so far untested therapeutic hypotheses. The diversity of methods, in many cases leveraging modern neural network architectures, reflects an intensely dynamic and explorative community, but despite the current hype surrounding AI-based drug discovery, a breakthrough in the field remains to be seen.

## METHODS

### Computational workflows

Computational methods are available from https://cache-challenge.org/results-cache-challenge-1

### Protein expression and purification

DNA fragments encoding LRRK2 residues (T2124-E2527) and (T2141-E2527) were cloned into pFastBac HTA donor plasmid downstream of a His-tag or into pFBD-BirA expression vector, a derivative of Invitrogen pFastBac Dual vector for in-cell biotinylation (https://www.thesgc-dev.org/sites/default/files/toronto_vectors/pFB-BirA.pdf), respectively. The resulting plasmid was transformed into DH10Bac™ Competent E. coli (Invitrogen) to obtain recombinant viral bacmid DNA, followed by a baculovirus generation for protein production in Sf9 insect cells. For in-cell biotinylation, D-biotin was added at the final concentration of 10 μg/mL during protein expression. The cells were harvested by centrifugation (2500 rpm for 10 mins at 10°C), 72-96 hours post-infection with well-developed signs of infections and 70-80 % viability as previously described.^27^ Harvested cells were resuspended in 20mM Tris-HCl, pH 7.5, 500mM NaCl, 5mM imidazole and 5% glycerol, 1X protease inhibitor cocktail (100 X protease inhibitor stock in 70% ethanol (0.25mg/ml Aprotinin, 0.25mg/ml Leupeptin, 0.25mg/ml Pepstatin A and 0.25mg/ml E-64) or Pierce™ Protease Inhibitor Mini Tablets, EDTA-free. The cells were lysed chemically by addition of 1mM PMSF, 1mM TCEP, 0.5% NP40 and benzonase (in-house) followed by sonication at frequency of 7.0 (5” on/7” off) for 5 min (Sonicator 3000, Misoni). The crude extract was clarified by high-speed centrifugation (60 min at 14000 rpm at 10°C) by Beckman Coulter centrifuge. The clarified lysate was loaded onto open columns containing pre-equilibrated Ni-NTA resin (Sigma Aldrich). The column was washed and eluted by running 20mM Tris-HCl, pH 7.5, 500mM NaCl, 5% glycerol, containing 5mM, 15mM and 250mM imidazole, respectively. The eluted proteins were then supplemented with 2mM TCEP. The His- and Avi-tagged protein was then further purified by size-exclusion chromatography on a Superdex200 16/600 using an ÄKTA Pure (Cytiva) after the column was equilibrated with 50mM Tris-HCl pH 7.5, 300mM NaCl, 2mM TCEP.

For the His-tagged protein, the tag was cleaved after elution using tobacco etch virus protease (TEV) overnight while the protein was dialyzed against 20mM Tris-HCl, pH 7.4, containing 300mM NaCl, 2mM TCEP. The protein was then loaded on equilibrated Ni-NTA resin for reverse affinity to remove His-tagged TEV enzyme and the uncut His-tagged proteins. The purity and size of the cut protein was confirmed on SDS-PAGE gel and mass spectrometry, respectively and the pure protein was concentrated and flash frozen.

### Surface plasmon resonance

The binding affinity of compounds was assessed by Surface plasmon resonance (SPR, Biacore™ 8K, Cytiva Inc.) at 25 °C. Biotinylated LRRK2 (2141-2527aa - https://www.addgene.org/210899/) was captured onto flow cells of a streptavidin-conjugated SA chip at approximately 5,000 response units (RU) (according to manufacturer’s protocol). Compounds were dissolved in 100% DMSO (30 mM stock) and diluted to 10 mM before serial dilutions were prepared in 100% DMSO (dilution factor of 0.33 was used to yield 5 concentrations). For SPR analysis, serially titrated compound was diluted 1:50 in HBS–buffer (10 mM HEPES pH 7.4, 150 mM NaCl, 0.01% Tween-20) to a final concentration of 2% DMSO. Experiments were performed using the same buffer containing 2% DMSO and multi-cycle kinetics with a 60 s contact time and a dissociation time of 120 s at a flow rate of 40 μL/min. Kinetic curve fittings and K_D_ value calculations were done with a 1:1 binding model using the Biacore Insight Evaluation Software (Cytiva Inc).

### Differential scanning fluorimetry

LRRK2 was diluted to 0.1 mg/mL in buffer (100 mM Hepes, 100 mM NaCl, pH 7.5) in the presence of 5x SYPRO Orange dye (Life Technologies, S-6650) and serially titrated compounds (up to 200 μM) in a total volume of 20 μL in a white polypropylene 384-well plate (Axygen, PCR-384-LC480-W). DSF was performed in a LightCycler 480 II (Roche Applied Science, Penzberg, Germany) using a 4°C/min temperature gradient from 20°C to 95°C. Data points were collected at 0.5°C intervals. DSF data was fitted to a Boltzmann sigmoid function and T_m_ values were determined as previously described^28^.

### Dynamic light scattering

The solubility of compounds was estimated by DLS that directly measures compound aggregates and laser power in solution. Compounds were serially diluted directly from DMSO stocks, then diluted 50x into filtered 10 mM HEPES pH 7.4, 150 mM NaCl(2% DMSO final). The resulting samples were then distributed into 384-well plates (black with a clear bottom, Corning 3540), with 20μl in each well. The sample plate was centrifuged at 3500 rpm for 5 minutes before loading into DynaPro DLS Plate Reader III (Wyatt Technology) and analyzed as previously described^29,30^.

### ^19^F-NMR spectroscopy

The binding of fluorinated compounds was assayed by looking for the broadening and/or perturbation of ^19^F resonances upon addition of LRRK2 (at protein to compound ratios of 0.5:1 to 4:1) in PBS buffer (pH 7.4, 137 mM NaCl, 2.7 mM KCl, 10 mM Na_2_HPO_4_, 1.8 mM KH_2_PO_4_, and with 5% D_2_O). 1D-^19^F spectra were collected at 298K on a Bruker AvanceIII spectrometer, operating at 600 MHz, and equipped with a QCI probe. Two to four thousand transients were collected with an acquisition period of 0.2 s, over a sweep width of 150 ppm, a relaxation delay of 1.5 s, and using 90° pulses centered at -120 ppm. The concentration of the compounds in both reference and protein-compound mixtures was 5-10 μM. TFA (20 μM) was added as an internal standard for referencing. Prior to Fourier transformation, an exponential window function was applied (lb = 1 to 3) to the FID. All processing was performed at the workstation using the software Topspin 3.5.

### Crystallization and structural determination

Human LRRK2 WDR domain (residues 2142–2527) was expressed, purified and crystallized as described previously (PMID: 30635421). Apo-LRRK2 WDR domain crystals were obtained by mixing equimolar amounts of protein (concentrated at 9 mg/mL) and precipitant solution containing 0.1 M Tris-HCl at pH 8.5, 1 M LiCl, 14% (w/v) polyethylene glycol (PEG) 6000, and 10% galactose in a manual plate vapour-diffusion hanging drops. LRRK2 crystals were then soaked into a 1 μL reservoir solution supplemented with 1 mM CACHE 1193-26 (dissolved from a previously prepared 100 mM DMSO stock solution) and 10% (v/v) Ethylene glycol for 2 hours at room temperature, then mounted and cryo-cooled in liquid nitrogen. Diffraction data were collected at the 24ID-E beamline at the Advanced Photon Source (APS). Dataset was processed with HKL3000^31^. Initial phases were obtained by using Apo-LRRK2 WDR domain (PDB ID:6DLO) as initial model in Fourier transform with refmac5^32^. Model building was performed in COOT^33^ and the structure was validated with Molprobity^34^. CACHE 1193-26 structure restraints were generated using Grade Web Server (http://grade.globalphasing.org).

## Supporting information

Supplemental Tables 1-8, including review, compound, and computation information

## Data availability

The crystal structure of LRRK2-WDR in complex with CACHE_1193_26 was deposited in the Protein Data Bank, PDB code 9C61. A generic description of all computational methods is available at https://cache-challenge.org/

## Acknowledgments

We thank Claudia Gordijo, Maxwell Morgan, and Richard Gold who played instrumental roles towards the successful launch of the CACHE challenges, and Shabbir Ahmad and Stuart Green for technical discussions on SPR assays. We thank Dr. Hao Wu for sharing the LRRK2 WDR domain construct for structural studies. We thank the staff at the Northeastern Collaborative Access Team, which is funded by the National Institute of General Medical Sciences from the National Institutes of Health (P30 GM124165). Experimental testing was supported by an Open Science Drug Discovery grant from Canada’s Strategic Innovation Fund (SIF Stream 5) administered by Conscience and the Michael J Fox Foundation, and conducted at the Structural Genomics Consortium, a registered charity (no: 1097737) that receives funds from Bayer AG, Boehringer Ingelheim, Bristol Myers Squibb, Genentech, Genome Canada through Ontario Genomics Institute [OGI-196], Janssen, Merck KGaA (aka EMD in Canada and US), Pfizer and Takeda. This project has received funding from the Innovative Medicines Initiative 2 Joint Undertaking (JU) under grant agreement No 875510. The JU receives support from the European Union’s Horizon 2020 research and innovation programme and EFPIA and Ontario Institute for Cancer Research, Royal Institution for the Advancement of Learning McGill University, Kungliga Tekniska Hoegskolan, Diamond Light Source Limited. This communication reflects the views of the authors and the JU is not liable for any use that may be made of the information contained herein. The Eiger 16M detector on the 24-ID-E beamline is funded by an NIH-ORIP HEI grant (S10OD021527). This research used resources of the Advanced Photon Source, a U.S. Department of Energy (DOE) Office of Science user facility operated for the DOE Office of Science by Argonne National Laboratory under contract no. DE-AC02-06CH11357.

